# ePlant in 2021: New Species, Viewers, Data Sets, and Widgets

**DOI:** 10.1101/2021.04.28.441805

**Authors:** Ben Waese-Perlman, Asher Pasha, Chantal Ho, Amirahmad Azhieh, Yushan Liu, Alexander Sullivan, Vincent Lau, Eddi Esteban, Jamie Waese, George Ly, Cornelia Hooper, S. Evan Staton, Nicholas Brereton, Cuong Le, Rex Nelson, Shelley Lumba, David Goodstein, A. Harvey Millar, Isobel Parkin, Lewis Lukens, Juergen Ehlting, Loren Rieseberg, Frédéric Pitre, Anne Brown, Nicholas J. Provart

**Affiliations:** Department of Cell & Systems Biology / Centre for the Analysis of Genome Evolution and Function, University of Toronto, CANADA; Phytozome, Joint Genome Institute, Lawrence Berkeley National Laboratory, U.S.A; School of Molecular Sciences, ARC Centre for Plant Energy Biology – University of Western Australia, AUSTRALIA; Agriculture and Agrifood Canada / University of Saskatchewan, CANADA; Department of Plant Agriculture, University of Guelph, CANADA; Department of Biology, University of Victoria, CANADA; Department of Botany, University of British Columbia, CANADA; Department of Biological Sciences, Université de Montréal, CANADA; United States Department of Agriculture, U.S.A

## Abstract

ePlant was introduced in 2017 for exploring large *Arabidopsis thaliana* data sets from the kilometre to nanometre scales. In the past four years we have used the ePlant framework to develop ePlants for 15 agronomically-important species: maize, poplar, tomato, *Camelina sativa*, soybean, potato, barley, *Medicago truncatula*, eucalyptus, rice, willow, sunflower, *Cannabis sativa*, wheat and sugarcane. We also updated the interface to improve performance and accessibility, and added two new views to the Arabidopsis ePlant – the Navigator and Pathways viewers. The former shows phylogenetic relationships between homologs in other species and their expression pattern similarities, with links to view data for those genes in the respective ePlants. The latter shows Plant Reactome metabolic reactions. We also describe new Arabidopsis data sets including single cell RNA-seq data from roots, and how to embed ePlant eFP expression pictographs into any web page.

## Introduction

Vast amounts of biological data have been generated over the past 15 years. Traditional methods of analyzing data in multiple formats can add a layer of friction to the creative processes of hypothesis generation. We introduced ePlant four years ago to help plant biologists explore and make connections between large *Arabidopsis thaliana* data sets for any given gene/gene product (Waese et al., 2017). It includes views for exploring expression differences in ecotypes at a kilometre scale, expression levels in tissues and organs at the centimetre scale, sub-cellular localization at the millimetre scale, protein-DNA and protein-protein interactions at the micrometre scale and, finally, the tertiary structure of the molecule itself at the nanometre scale. Combining multiple views into one visual analytic platform allows researchers to ask and answer complex biological questions about a gene of interest using a single, user-friendly interface.

Development of ePlant has continued since the original paper was published. Several new views have been added and the user interface has been updated to improve performance and accessibility. The new views include a Navigator viewer, a Pathways viewer, and seven new eFP viewers based on RNA-seq data. In addition, 15 new ePlants have been developed for agronomically-important species beyond *Arabidopsis thaliana*. Last, we have made it easy to embed ePlant eFP images showing expression patterns into any web page as a widget.

## Results

### New ePlant Design with Improved Accessibility and Performance

Since the original Waese et al. (2017) paper, ePlant interface has undergone a major update (see **Figure 1**). The colours have been updated to improve contrast, infrequently used features were removed to reduce clutter, and animated page transitions were removed to eliminate negative experiences that motion effects cause for some users. In addition, gene loading times have improved by up to 300% and several bugs have been fixed that used to cause unpredictable behavior when asynchronous events did not happen as expected.

**Figure 1:**
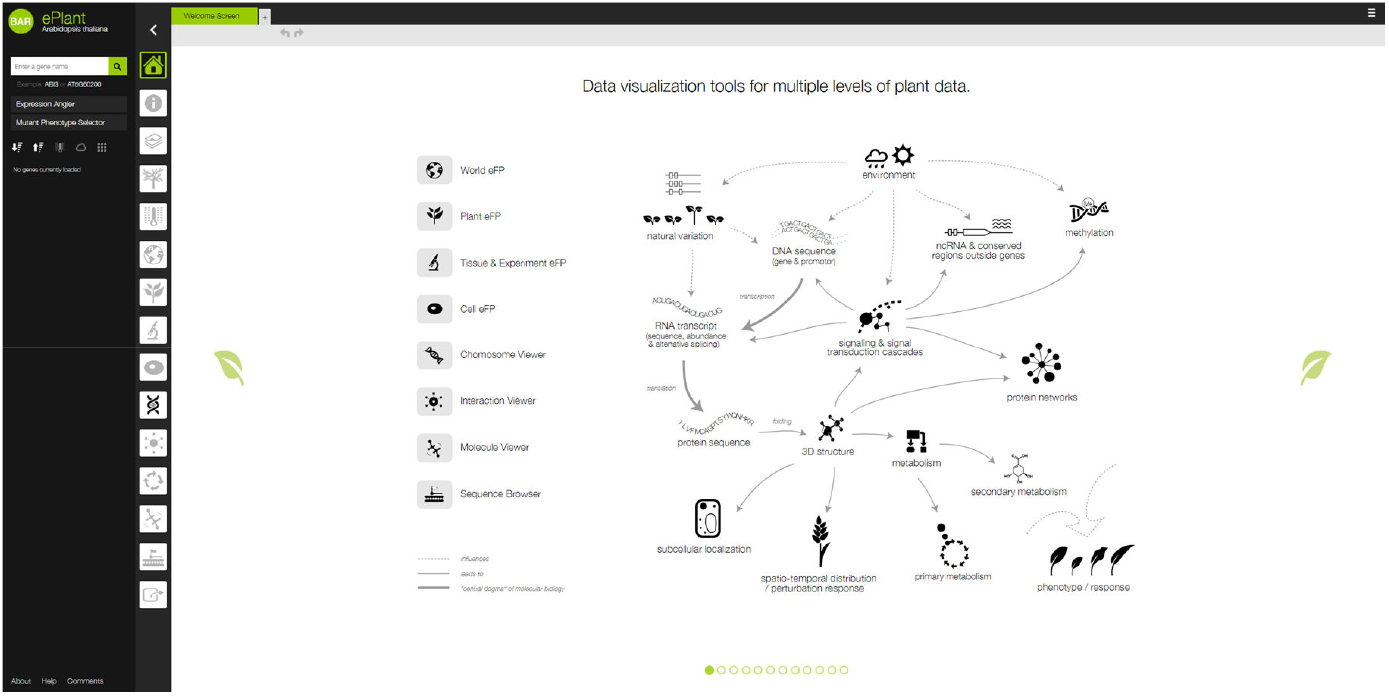
The new ePlant home page.

To improve contrast, we swapped the background colours for darker ones, removed gradients and changed text colours, making much of ePlant compatible with the higher WCAG “AAA” standard for enhanced contrast, which requires a ratio of at least 7:1^1^. The original design did not meet the WCAG “AA” standard. The new design uses the colours shown in **Figure 2**.

**Figure 2:**
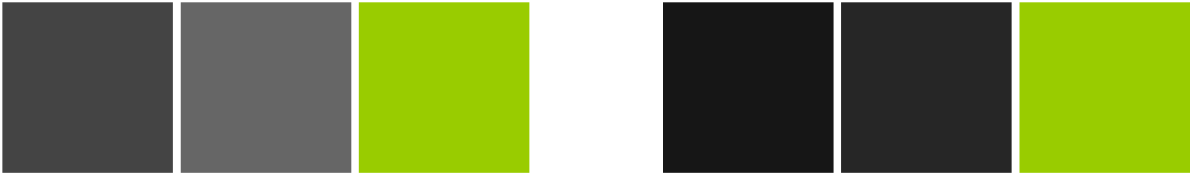
Original ePlant colours (left) #444444, #666666, #99CC00; new colours (right) #161616, #262626, #99CC00

We have maintained use of the “BAR green” as the highlight colour, despite darkening the background colours. By swapping the background colours to darker alternatives and removing the use of white text on a green background we were able to increase contrast significantly and create a more readable design (**Figure 3**). We also removed the animated transitions between views to address concerns from researchers with vestibular disorders^2^.

**Figure 3:**
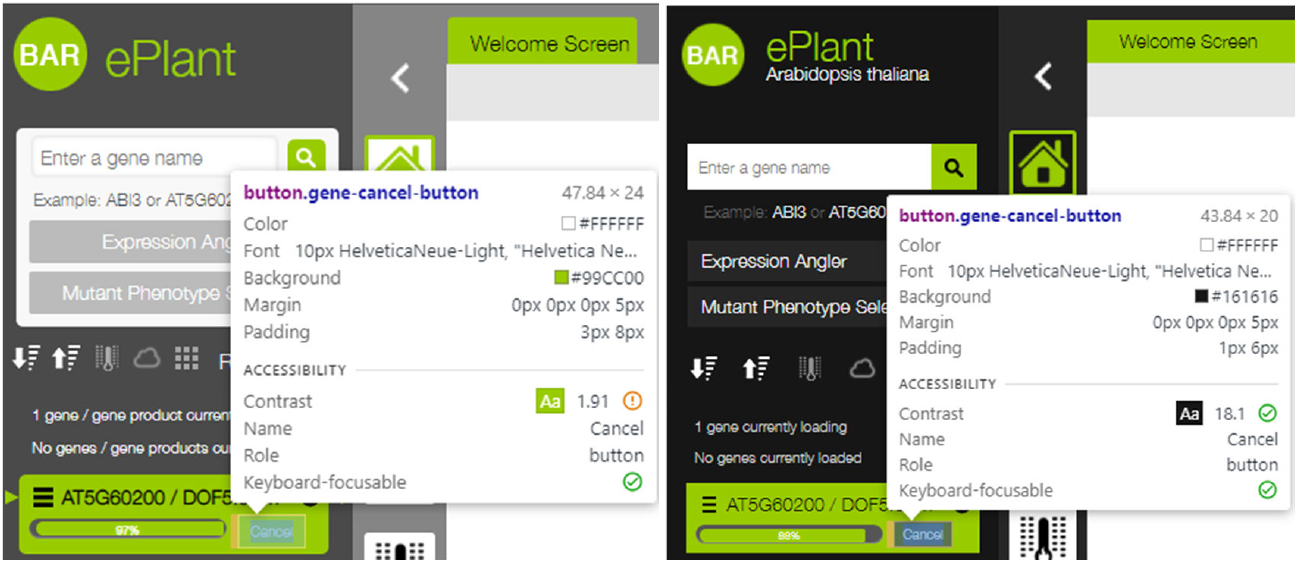
Contrast of the original ePlant design versus the new design. The original design (left) does not meet the WCAG “AA” level for contrast, while the new design (right) exceeds the “AAA” contrast level by a factor of more than two, as shown using Chrome’s developer tools.

### A Small Farm of ePlants

The original ePlant published by Waese et al. (2017) only supported data from *Arabidopsis thaliana*. We now have ePlants for another 15 species: *Arabidopsis thaliana*, maize, poplar, tomato, *Camelina sativa*, soybean, potato, barley, *Medicago truncatula*, eucalyptus, rice, willow, sunflower, *Cannabis sativa*,wheat and sugarcane (see **Figure 4** and **Table 1**).

**Figure 4:**
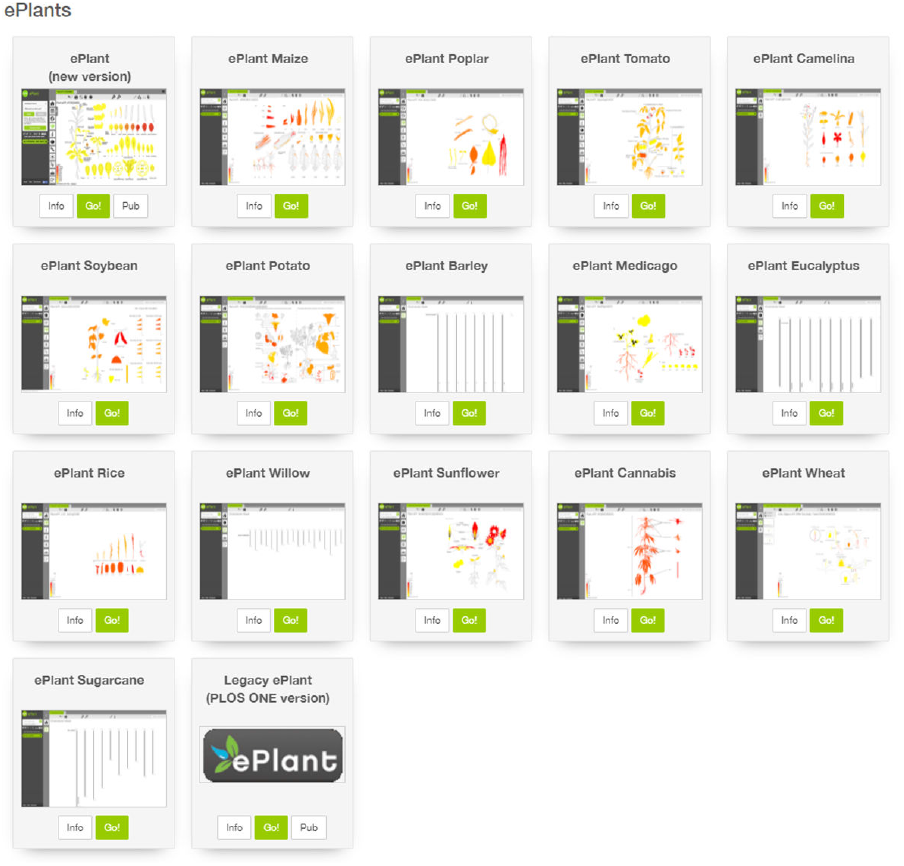
Accessing ePlants from the BAR homepage at http://bar.utoronto.ca.

**Table 1:**
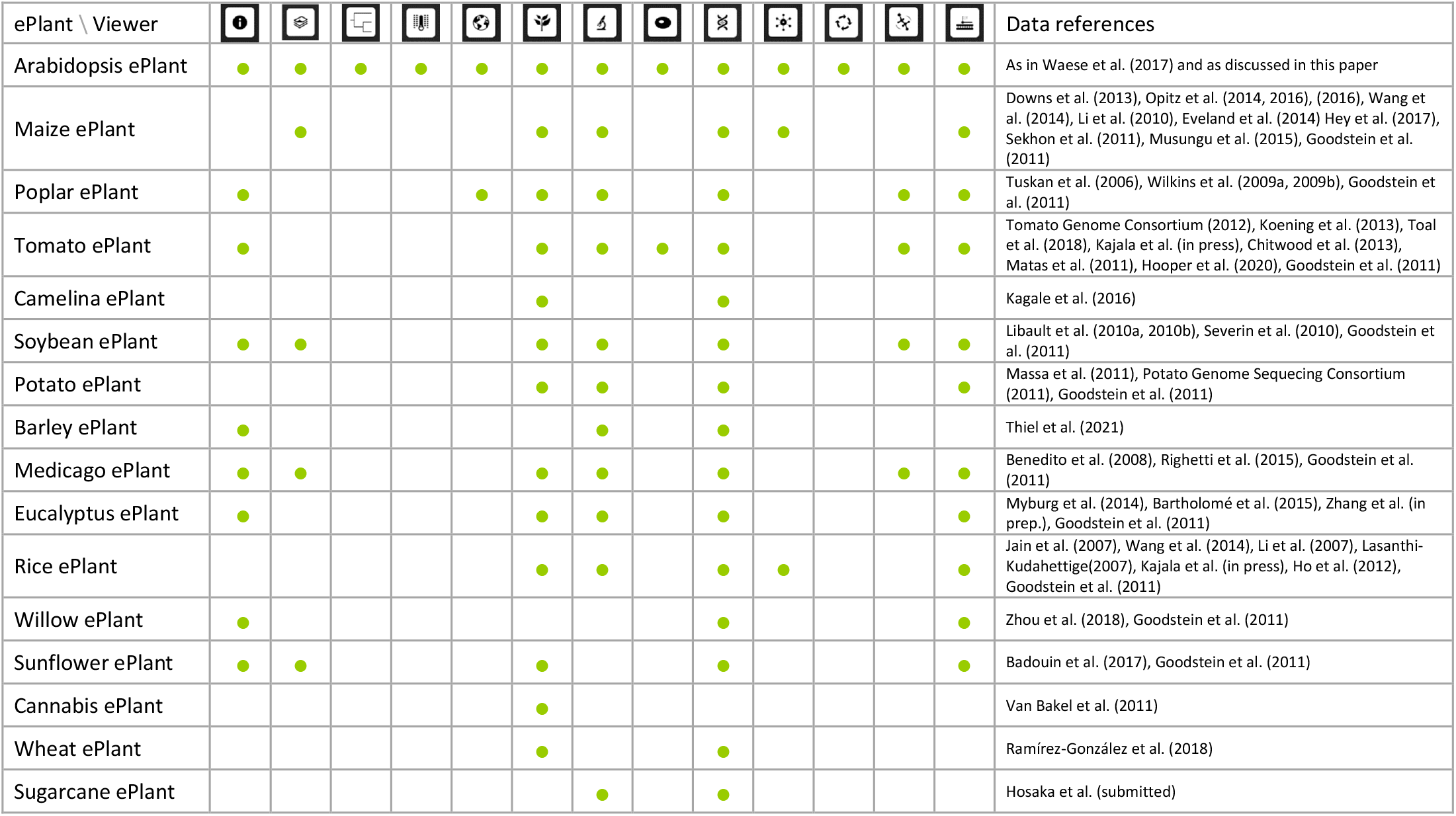
Overview of Data Available in each ePlant. Icons are defined in Figure 1, with the exception of the first 4: the 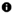 icon denotes gene information, the 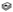 icon denotes publications, the 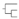 icon denotes ePlant Navigator, and the 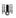 icon denotes the Heatmap Viewer. A new icon is the Pathways Viewer 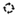 icon. ePlants are available at http://bar.utoronto.ca.

| ePlant \ Viewer |  |  |  |  |  |  |  |  |  |  |  |  |  | Data references |
| --- | --- | --- | --- | --- | --- | --- | --- | --- | --- | --- | --- | --- | --- | --- |
| Arabidopsis ePlant | ● | ● | ● | ● | ● | ● | ● | ● | ● | ● | ● | ● | ● | As in Waese et al. (2017) and as discussed in this paper |
| Maize ePlant |  | ● |  |  |  | ● | ● |  | ● | ● |  |  | ● | Downs et al. (2013), Opitz et al. (2014, 2016), (2016), Wang et al. (2014), Li et al. (2010), Eveland et al. (2014) Hey et al. (2017), Sekhon et al. (2011), Musungu et al. (2015), Goodstein et al. (2011) |
| Poplar ePlant | ● |  |  |  | ● | ● | ● |  | ● |  |  | ● | ● | Tuskan et al. (2006), Wilkins et al. (2009a, 2009b), Goodstein et al. (2011) |
| Tomato ePlant | ● |  |  |  |  | ● | ● | ● | ● |  |  | ● | ● | Tomato Genome Consortium (2012), Koenig et al. (2013), Toal et al. (2018), Kajala et al. (in press), Chitwood et al. (2013), Matas et al. (2011), Hooper et al. (2020), Goodstein et al. (2011) |
| Camelina ePlant |  |  |  |  |  | ● |  |  | ● |  |  |  |  | Kagale et al. (2016) |
| Soybean ePlant | ● | ● |  |  |  | ● | ● |  | ● |  |  | ● | ● | Libault et al. (2010a, 2010b), Severin et al. (2010), Goodstein et al. (2011) |
| Potato ePlant |  |  |  |  |  | ● | ● |  | ● |  |  |  | ● | Massa et al. (2011), Potato Genome Sequencing Consortium (2011), Goodstein et al. (2011) |
| Barley ePlant | ● |  |  |  |  |  | ● |  | ● |  |  |  |  | Thiel et al. (2021) |
| Medicago ePlant | ● | ● |  |  |  | ● | ● |  | ● |  |  | ● | ● | Benedito et al. (2008), Righetti et al. (2015), Goodstein et al. (2011) |
| Eucalyptus ePlant | ● |  |  |  |  | ● | ● |  | ● |  |  |  | ● | Myburg et al. (2014), Bartholomé et al. (2015), Zhang et al. (in prep.), Goodstein et al. (2011) |
| Rice ePlant |  |  |  |  |  | ● | ● |  | ● | ● |  |  | ● | Jain et al. (2007), Wang et al. (2014), Li et al. (2007), Lasanthi-Kudahettige(2007), Kajala et al. (in press), Ho et al. (2012), Goodstein et al. (2011) |
| Willow ePlant | ● |  |  |  |  |  |  |  | ● |  |  |  | ● | Zhou et al. (2018), Goodstein et al. (2011) |
| Sunflower ePlant | ● | ● |  |  |  | ● |  |  | ● |  |  |  | ● | Badouin et al. (2017), Goodstein et al. (2011) |
| Cannabis ePlant |  |  |  |  |  | ● |  |  |  |  |  |  |  | Van Bakel et al. (2011) |
| Wheat ePlant |  |  |  |  |  | ● |  |  | ● |  |  |  |  | Ramírez-González et al. (2018) |
| Sugarcane ePlant |  |  |  |  |  |  | ● |  | ● |  |  |  |  | Hosaka et al. (submitted) |

Although they share a common interface, the ePlants are not all identical. The 16 ePlants draw data from a variety of databases, and different species have different views based on data availability, as shown in **Table 1**. The Arabidopsis ePlant continues to have the greatest number of views. We developed a pipeline for creating a new ePlant of interest and this is available at https://github.com/BioAnalyticResource/ePlant_Pipeline. For several ePlants, we have predicted “structure-omes” using Phyre2 (Kelley et al., 2015).

### Two New Viewers for the Arabidopsis ePlant

#### Navigator Viewer

The Navigator Viewer as shown in **Figure 5** provides a tree-based visualization of homologous genes in other species. It uses data from the BAR’s Expressolog database (Patel et al., 2012) to construct a phylogenetic tree with leaf nodes representing the homologs from different species. For each gene, its species is reported along with the similarity at the sequence level and at the level of expression pattern similarity, based on the expressolog method developed by Patel et al. (2012). In this way, genes that are both sequence similar and have similar patterns of expression in equivalent tissues may be easily identified. Link-outs are also provided to each gene’s corresponding ePlant and or eFP view. In addition, dynamic links to genomevolution.org’s CoGe database (Lyons and Freeling, 2008) and to the Gramene database (Tello-Ruiz et al., 2018) are provided so that a researcher can easily access further information about syntenic relationships etc.

**Figure 5:**
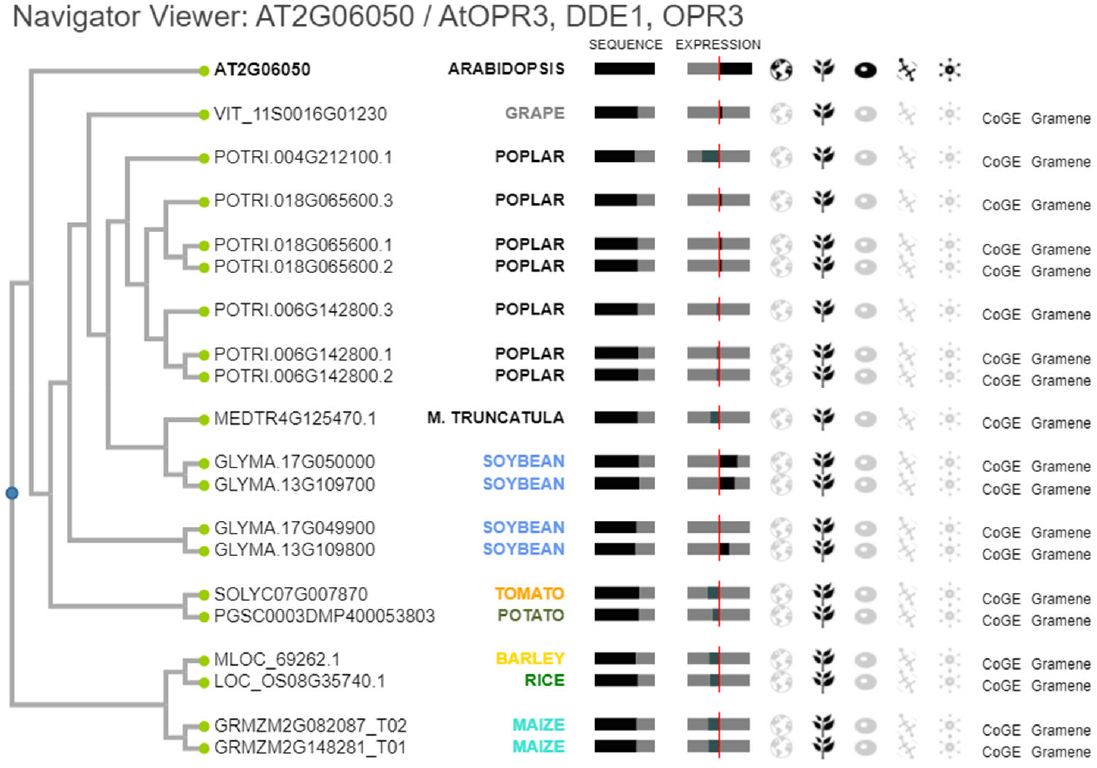
ePlant Navigator Viewer

This tool was created with the goal of quickly and easily helping biologists find similar genes across the various ePlant species. If a biologist studying one species is familiar with a gene in another, this view allows them to make a connection between the two.

#### Pathways Viewer

The Pathways Viewer shown in **Figure 6** displays metabolic pathways associated with the active gene. It draws data from the Plant Reactome pathway database (Naithani et al., 2019) via application programming interface (API) calls and generates a pathway diagram. This is done using Cytoscape.js, a JavaScript-based graph theory library for network visualization and analysis (Franz et al., 2016). Nodes are laid out with a force directed layout known as CoSE Bilkent^3^ and styled appropriately. All pathways available for a given gene are displayed as selectable tabs and a direct link to the pathway in the Plant Reactome pathway browser is provided for more information. In addition, hovering over the active gene label, which is denoted by bold text, calls up a tooltip which contains the Klepikova Plant eFP view for that gene, based on the Klepikova developmental atlas (Klepikova et al., 2016) – see the next section. The Klepikova eFP view can be used to assess in which parts of the plant the gene is strongly expressed, to help identify parts of the plant where the depicted reaction might be occurring.

**Figure 6:**
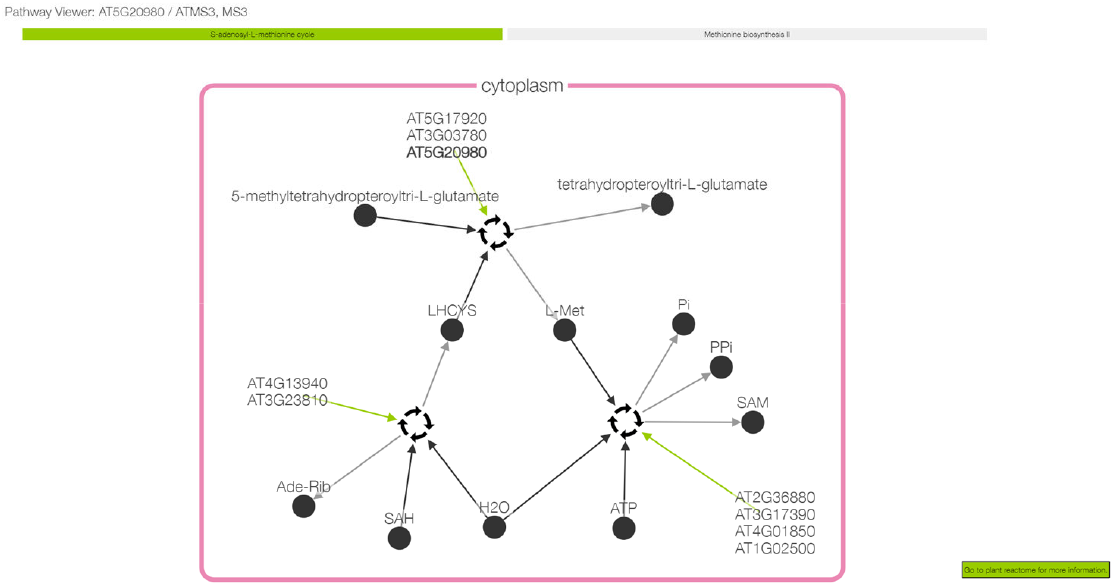
ePlant Pathways Viewer

The ePlant Pathways Viewer fills a gap in the conceptual hierarchy within ePlant, between the Interaction Viewer and the Molecule Viewer. Currently this view is only available for the Arabidopsis ePlant with data for over 236 pathways, 590 reactions and 1059 gene products and it is coming to the other ePlants soon. Plotting these pathways enables researchers to understand the biological context of the active gene product. The other views in ePlant show where in the cell the gene/macromolecule exists and how it is expressed, but this view is what makes it possible to answer the question “what is the gene doing in the cell” in a way that was not possible before.

### New Data Sets

The original ePlant publication introduced the Plant eFP Viewer and the Tissue and Experiment eFP Viewers with more than 20 different views. They display pictographs representing the level of expression of a queried gene in organs of the plant or in specific tissues or cell types, with intensity of expression represented by a colour gradient. Since the original publication in 2017, a number of new views have been added, with the Single Cell RNA Sequence eFP (Ryu et al., 2019) being one of six added in this new update, as shown in **Figure 8**. Most of the newly added data are based on RNA sequencing and include a new Plant eFP Viewer for the Klepikova RNA-seq-based developmental atlas shown in **Figure 7** (Klepikova et al., 2016), a Germination view (Narsai et al., 2011), a Shoot Apex view (Tian et al., 2014), a Root Immunity Elicitation view (Rich-Griffin et al., 2020), and a Guard Cell Drought view (van Weringh et al., 2021).

**Figure 7:**
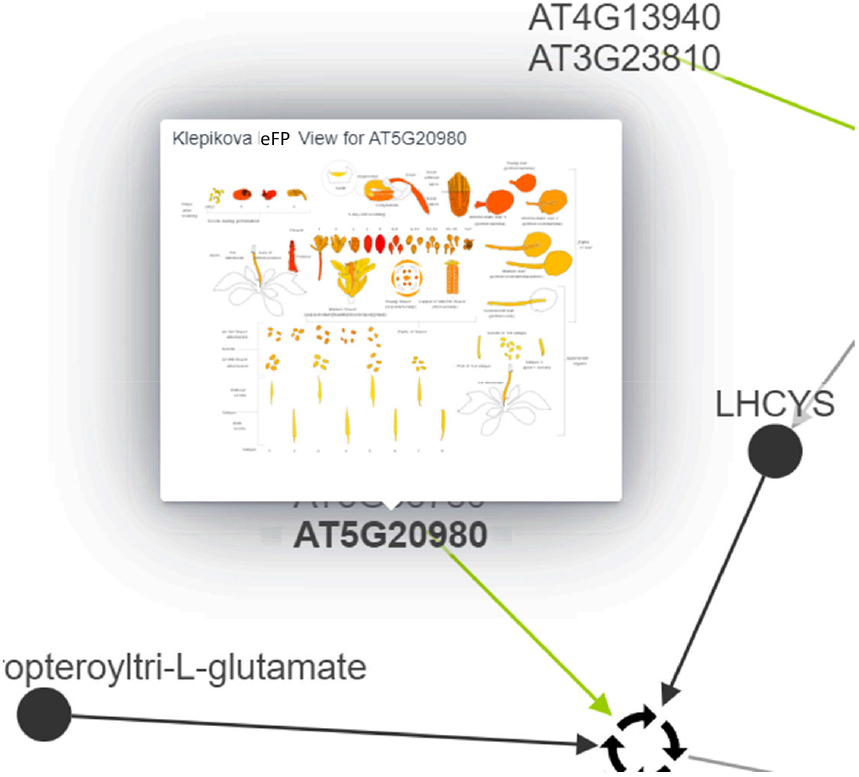
Klepikova Plant eFP tooltip in ePlant’s Pathways Viewer, showing the expression pattern of At5g20980, ATMS3, involved in the S-adenosyl-L-methionine cycle, showing strong expression (red colour) in young leaves, flower buds, stems, and germinating seeds.

**Figure 8:**
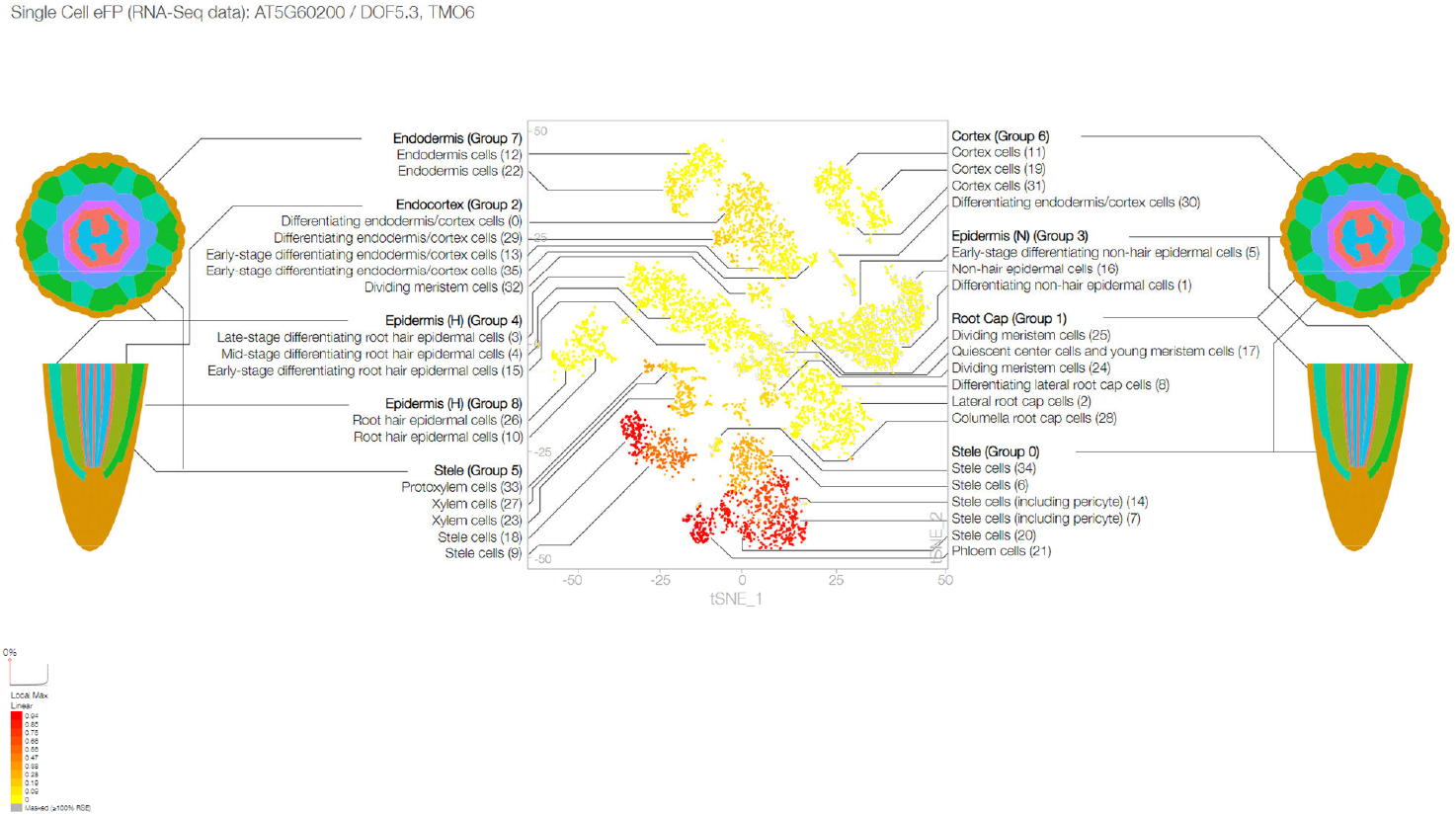
Single Cell RNA-seq eFP View for the gene At5g60200

We also updated the non-synonymous single nucleotide polymorphism data source for the Arabidopsis Molecule Viewer from the 1001 Proteomes site (Joshi et al., 2012), which had been deprecated, to the 1001 Genomes API (1001 Genomes Consortium, 2016). The advantage of doing this, apart from having a reliable data source, is that the 1001 Genomes API is more comprehensive, containing polymorphism data for almost twice as many Arabidopsis ecotypes.

### ePlant eFP Widgets

We have developed SVG-based ePlant Plant eFP widgets for use in any website, as shown in **Figure 9**. See https://bar.utoronto.ca/~asullivan/ePlant_Plant_eFP/example/ for how to use these. The widgets behave similarly to the eFP views found within ePlant itself and are available for 88 different SVGs. They function after the same paradigm, with a gradient showing intensity of gene expression in different tissue regions or cell types taken from different parts of the plant.

**Figure 9:**
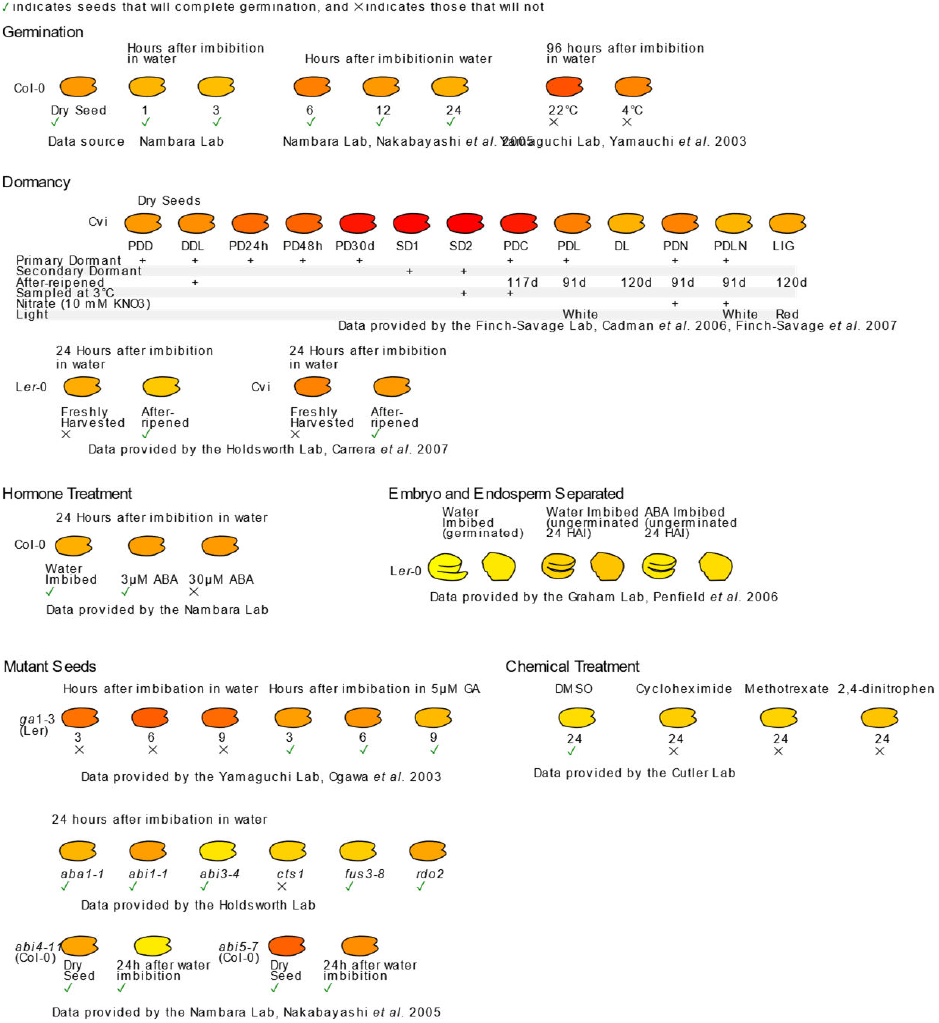
Seed ePlant eFP widget

## Discussion

We present several useful updates to our ePlant tool. Improvements to the accessibility and performance of the interface makes for a better user experience. Fifteen new ePlants permit easy exploration of expression, interaction, subcellular localization and structure data across multiple agronomically-important species. We hope that the research communities for these species will be interested in adding data sets to these new ePlants in the future. Two new viewers, the Navigator and Pathways viewers, permit genes/gene products to be explored in the context of their homologs and biochemical pathways retrieved from Plant Reactome, respectively. A Plant eFP tooltip for the active gene/gene product in the latter viewer can help highlight in which organ a pathway might be operating. We have added new expression data sets based on RNA sequencing to the Tissue and Experiment Viewers, increasing the breadth of transcriptomic data viewable in a pictographic manner. Last, ePlant eFP widgets may be easily embedded in any webpage for any Arabidopsis gene with some straightforward code.

## Acknowledgements, Contributions, and Funding

We are grateful to Pankaj Jaiswal and Justin Preece from Oregon State University for helpful discussions about using the Plant Reactome API. BWP, AA and YL implemented the Pathways Viewer. BWP reskinned the UI and made many improvements to the ePlant code base. AP databased additional data not already on the BAR and customized ePlant code for new ePlants, and developed APIs for use in the Pathways Viewer. C. Ho developed the ePlant Navigator. AS created the ePlant Widgets. JW, VL and NJP supervised students (BWP, CH, YL, AA), and assisted with image preparation for new ePlant views, as did EE. GL, SL, IP, LL, CL, RN, JE, LR, ES, FP and AB provided data, images, and feedback for new ePlants. AHM and C. Hooper provided data from CropPAL (http://crop-pal.org/). DG assisted with Phytozome JBrowse deployment in the ePlants. BWP, JW and NJP wrote and edited the manuscript and created the figures. NJP was funded by a grant from Genome Canada/Ontario Genomics (OG-128).

## Footnotes

1 https://www.w3.org/TR/WCAG21/#contrast-minimum

2 https://alistapart.com/article/designing-safer-web-animation-for-motion-sensitivity/

3 https://github.com/cytoscape/cytoscape.js-cose-bilkent

